# Examining the role of *Acinetobacter baumannii* Plasmid Types in Disseminating Antimicrobial Resistance

**DOI:** 10.1101/2023.09.03.556112

**Authors:** Margaret M. C. Lam, Mehrad Hamidian

**Affiliations:** Department of Infectious Diseases, Central Clinical School, Monash University, Melbourne, VIC, Australia; Australian Institute for Microbiology & Infection, University of Technology Sydney, Ultimo, NSW, Australia

**Keywords:** *Acinetobacter baumannii*, antimicrobial resistance gene, AMR, plasmid, plasmid replication, Rep, *rep*, sequence type, and carbapenem resistance

## Abstract

*Acinetobacter baumannii* is a Gram-negative pathogen responsible for hospital-acquired infections with high levels of antimicrobial resistance (AMR). The spread of multidrug-resistant *A. baumannii* strains, particularly those resistant to carbapenems, has become a global concern. Spread of AMR in *A. baumannii* is primarily mediated by the acquisition of AMR genes through mobile genetic elements, such as plasmids. Thus, a comprehensive understanding of the role of different plasmid types in disseminating AMR genes is essential. In this study, we analysed the distribution of plasmid types, sampling sources, geographic locations, and AMR genes carried on *A. baumannii* plasmids. A collection of 814 complete plasmid entries was collated and analysed. Most plasmids were identified in clinical isolates from East Asia, North America, South Asia, West Europe, and Australia.

We previously devised an *Acinetobacter* Plasmid Typing (APT) scheme where *rep/*Rep types were defined using 95% nucleotide identity and updated the scheme in this study by adding 13 novel *rep*/Rep types (93 types total). The APT scheme now includes 178 Rep variants belonging to three families: R1, R3, and RP. R1-type plasmids were mainly associated with global clone 1 strains, while R3-type plasmids were highly diverse and carried a variety of AMR determinants including carbapenem, aminoglycoside and colistin resistance genes. Similarly, RP-type and rep-less plasmids were also identified as important carriers of aminoglycoside and carbapenem resistance genes. This study provides a comprehensive overview of the distribution and characteristics of *A. baumannii* plasmids, shedding light on their role in the dissemination of AMR genes. The updated APT scheme and novel findings enhance our understanding of the molecular epidemiology of *A. baumannii* and provide valuable insights for surveillance and control strategies.

**IMPORTANCE:** *A. baumannii* has emerged as a major cause of nosocomial infections, particularly in intensive care units, posing a substantial challenge to patient safety and healthcare systems. Plasmids, which carry antimicrobial resistance (AMR) genes, play a crucial role in the multidrug resistance exhibited by *A. baumannii* strains, necessitating a comprehensive understanding of plasmid spread, and how to track them. This study provides important insights into *A. baumannii* plasmid epidemiology, and the extent of their role in spreading clinically significant AMR genes and how they are differentially distributed across different clones i.e. sequence types (STs) and geographical regions. These insights are important for identifying high-risk areas or clones implicated in plasmid transmission, in the context of the spread of multidrug-resistant *A. baumannii* strains. It also highlights the involvement of R3-type, RP-type and rep-less plasmids in the acquisition and spread of significant AMR genes including those conferring resistance to carbapenems, aminoglycosides and colistin.

## INTRODUCTION

*Acinetobacter baumannii* is a notorious opportunistic Gram-negative pathogen that causes hospital-acquired infections, such as bacteraemia, pneumonia, wound infections and urinary tract infections ^1^ that are difficult — and in many cases impossible — to treat due to high levels of resistance to several antimicrobial classes ^2,3^. Indeed, the spread of *A. baumannii* strains resistant to all available antimicrobials has become a major global concern ^3,4^, and carbapenem-resistant *A. baumannii* has been flagged by the World Health Organisation as the number one priority for antimicrobial development ^5^. In *A. baumannii*, antimicrobial resistance (AMR) is known to occur primarily by the horizontal acquisition of AMR genes via mobile genetic elements (MGEs) such as transposons and plasmids ^3,6–8^. Notably, *A. baumannii* plasmids are increasingly recognised as a major source for disseminating AMR genes such as those that confer resistance to carbapenems and colistin ^9–13^.

*A. baumannii* has a unique repertoire of plasmids that capture and mobilise a wide range of genetic material involved in pathogenesis and AMR (6, 15-18). Recently, we developed a plasmid typing scheme based on the sequence of replication initiation genes from 621 complete *A. baumannii* plasmids called *Acinetobacter* Plasmid Typing (APT) scheme ^13^. This first version of the APT scheme includes 80 Rep types belonging to three families; R1 types 1 to 6, R3 types 1 to 69, and RP types 1-5 (https://github.com/MehradHamidian/AcinetobacterPlasmidTyping) ^13^. However, the role of each plasmid type in dissemination of AMR genes remains to be established. To date, several studies have examined the role of *A. baumannii* plasmids in the spread of AMR genes ^9,10,14–17^ but most of these studies have only reported on individual or a limited number of plasmids and thus, a comprehensive overview of how various plasmid types are involved in the dissemination of AMR genes remains elusive. In this study, we address this gap by examining the distribution of chromosomal sequence types, sampling, geographies, and AMR genes carried on plasmids originally included in the APT database alongside an additional 193 complete plasmids that have since been deposited in GenBank (as of August 18, 2022). We also provide an update to the original APT scheme with the addition of novel *rep*/Rep types.

## RESULTS AND DISCUSSION

### Overview of genome and plasmid dataset

As of August 18, 2022, 450 complete *A. baumannii* genomes were available in GenBank. Of these 450 complete genomes, 80% (n=355) had at least one plasmid (Table S1) with 236 genomes containing one (n=1) plasmid, and 113 carrying two plasmids (Table 1 and Table S1). Ninety-one (n=95) genomes lacked a plasmid and were not studied here. To broaden our plasmid dataset, we extended our search to the RefSeq database and captured an additional 92 genomes that contained at least one (n=1) plasmids. Of these 92 genomes/unique strains, n=63 were not linked to a genome project and n=29 genomes were sourced from WGS (Whole Genome Shotgun), which included draft genomes with complete plasmid entries. Following curation of the dataset (i.e. exclusion of duplicate entries and assembly QC; see methods for more details), our final dataset was comprised of 814 non-redundant plasmid entries corresponding to at least 440 unique isolates (Table S1 and Table S2; n=2/814 plasmids unassigned due to absence of BioSample and strain name). Notably, of the 440 unique genomes/isolates, the *rep*/Rep sequences of n=329 were had already been analysed in the original APT scheme ^13^. Indeed, in this study, we investigated the prevalence of antimicrobial resistance (AMR) genes within different plasmid types, considering their associated meta-data. A total of 440 unique genomes/strains were analysed, comprising both the 329 genomes previously reported and an additional 111 genomes captured in this study. Over half of the isolates carried one plasmid (n=229 isolates; 52%) and 27% (n=120) carried two plasmids (Table 1). Seven genomes contained 6-11 plasmids, indicating that some *A. baumannii* strains have the capacity to carry a significantly high load of plasmids.

**TABLE 1.** Number of plasmids found in 440 unique BioSamples and/or strain names.

| No. of plasmids<br>per genomes | No. of<br>genomes |
| --- | --- |
| 1 | 229 (53%) |
| 2 | 120 (25%) |
| 3 | 52 (10%) |
| 4 | 28 (7%) |
| 5 | 4 (2%) |
| 6 | 3 (0.2%) |
| 8 | 2 (0.2%) |
| 9 | 1 (0.12%) |
| 11 | 1 (0.2%) |

Of the 440 unique isolates, ST2 represented the most abundant sequence type (n=199), followed by ST1 (n=25), ST25 (n=14), and ST622 (n=10). In 2019, we reported that both the geographies and sampling types of publicly available *A. baumannii* genomes deposited in NCBI were extremely skewed ^3^. This is similarly reflected in this dataset, with the vast majority sequenced in strains recovered from clinical samples (n=359; 81.5%), and few from non-clinical sources (FIG 1). This is primarily because of continued significance and attention paid towards clinical strains, and a lessened focus on studying the role of non-clinical reservoirs (e.g. environment, animals) as potential sources for plasmids and AMR genes. Most complete plasmid entries were sequenced from isolates collected in East Asia (primarily China, n=105/155 isolates), followed by North America (primarily US, n=77/90) and South Asia (primarily India, n=44/46). The strains were isolated from forty-three countries, and those with at least ten plasmids plus isolates within the dataset included South Korea (n=38), Australia (n=15), France (n=15), Canada (n=13), Iraq (n=12), Mexico (n=12), and Italy (n=10; see FIG 1 and Table S1 and Table S2).

**FIG 1.**
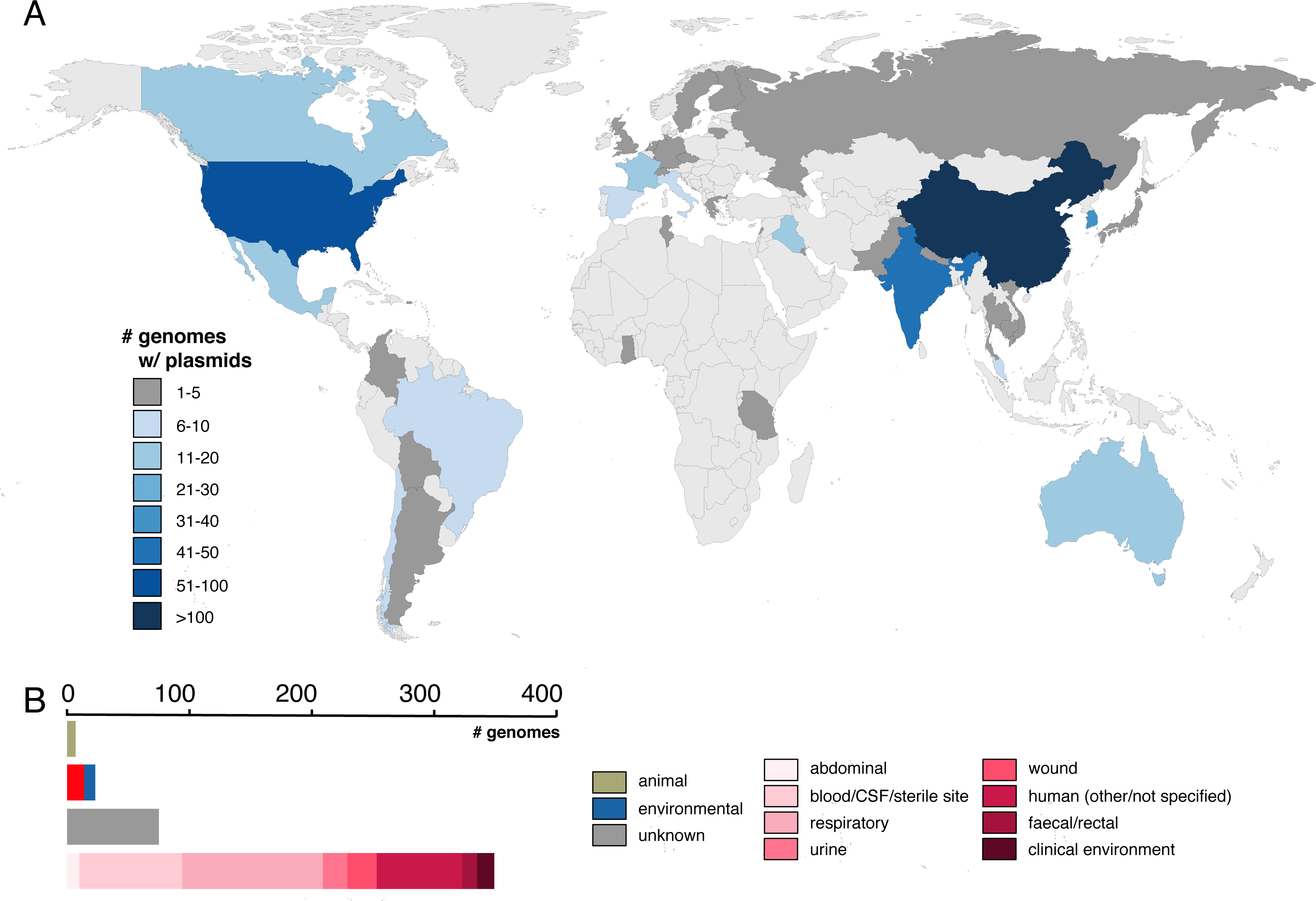
Geographical distribution and isolation sources of publicly available *A. baumannii* strains carrying plasmids. A) Geographical distribution of plasmid-containing strains, colour-coded by the number of genomes accessible in GenBank as of August 2022. B) Isolation sources of plasmid-carrying strains, depicted with a scale bar provided above.

### Novel *rep* sequences and update to the *Acinetobacter* Plasmid Typing (APT) scheme

Of the 814 plasmids studied here, 621 (i.e. 71%) were previously used to generate the first version of the APT scheme. An additional 193 complete plasmid entries, corresponding to 92 isolates, had since been released in GenBank between February 2021 and mid-August 2022. Using the criteria we had previously established for assigning *rep* types (i.e., 95% DNA identity) ^13^, n=13 novel *rep* types were identified (n=10 R3 types, designated R3-T70 to R3-T79; n=2 R1 types, R1-T7 and R1-T8; n=1 RP type, RP-T6), resulting in a total of n=78, 7 and 6 R3, R1 and RP types, respectively (Table 2, Table S2). In addition to the novel *rep*/Rep sequences, we also report additional updates to the scheme as follows. R3-T49 has been removed from the updated APT scheme, as the corresponding *rep* sequence (previously r3-T49_NZ_AYFZ01000080.1_pABUH2a-5.6_c33) has been identified as a R3-T26 variant. Specifically, this variant carries an insert of 84 bp that differentiates this sequence from the other R3-T26 variants. This entry has been subsequently renamed to R3-T26* and R3-T49 retired from the scheme. R1-T3 was also retired due to possible sequencing/assembly errors resulting in shortening the Rep reading frame by approximately 150 amino acids. Lastly, we highlight two corrections to Figure 3 and Table 3 published in the original APT paper ^13^: i) R3-T3 was annotated twice in FIG 3; the first annotated clade highlighted in orange should be corrected to R3-T8, and ii) in Table 3, the rows and columns should read R1-T1 to R1-T6 (i.e. not P1-T1 to P1-T6).

**TABLE 2.** Number of plasmids in each *rep* family.

| Rep plasmid type/<br>family | Pfam | No. of<br>plasmids | No. of plasmids<br>with AMR | *rep* types<br>(95% identity) |
| --- | --- | --- | --- | --- |
| R1 (Rep_1) | 01446 | 16 | 0 | 7 |
| R3 (Rep_3) | 01051 | 479 | 122 (25.5%) | 78 |
| RP (RepPriCT_1) | 03090 | 158 | 64 (40.5%) | 6 |
| No Rep | NA | 161 | 109 (67.7%) | NA |
| **Total** | NA | **814** | **295 (36.2%)** | **92** |

**TABLE 3.** Distribution of plasmids corresponding to the 15 most abundant R3 Rep types in major globally distributed sequence types (STs)

| R3-Types | Total | AMR | GC1 | GC2 | ST10 | ST15 | ST17 | ST25 | ST79 | ST85 | ST103 | ST622 | ST649 | Other<br>STs | Novel<br>STs |
| --- | --- | --- | --- | --- | --- | --- | --- | --- | --- | --- | --- | --- | --- | --- | --- |
| R3-T1 | 116 | **24** | **21** | **56** | 2 | - | - | 4 | 1 | - | - | 8 | - | 15 | 9 |
| R3-T3 | 75 | 9 | - | **51** | 1 | - | - | 1 | - | 1 | - | - | - | 21 | - |
| R3-T2 | 33 | 8 | 1 | **15** | - | - | - | 1 | - | - | - | - | 1 | 9 | 6 |
| R3-T4 | 33 | 1 | 1 | **23** | - | - | - | - | - | - | - | - | - | 9 | - |
| R3-T5 | 14 | 1 | - | 1 | 3 | - | - | - | 1 | - | - | - | 1 | 8 | - |
| R3-T14 | 14 | 9 | 1 | 1 | - | 4 | - | - | 2 | - | - | - | - | 4 | 2 |
| R3-T7 | 13 | 4 | 2 | 1 | - | - | - | - | - | - | - | - | - | 9 | 1 |
| R3-T13 | 12 | 3 | - | - | - | 1 | - | 1 | - | - | - | - | - | 10 | - |
| R3-T6 | 11 | 1 | - | - | - | - | - | - | 2 | 2 | - | - | - | 6 | 1 |
| R3-T15 | 10 | 0 | - | 2 | - | - | - | 2 | - | - | - | - | 1 | 4 | 1 |
| R3-T8 | 9 | 7 | - | 4 | - | - | - | - | - | - | - | - | - | 1 | 4 |
| R3-T10 | 8 | 3 | 1 | 1 | - | - | 1 | - | - | - | 2 | - | - | 3 | - |
| R3-T11 | 8 | 0 | 1 | 2 | 2 | - | - | - | - | - | - | - | - | 3 | - |
| R3-T9 | 8 | 4 | - | **8** | - | - | - | - | - | - | - | - | - | - | - |
| R3-T12 | 7 | 2 | - | - | - | - | - | 1 | - | - | - | - | - | 6 | - |

### Plasmids encoding the Rep_1 family replication protein (Rep_1 or R1 plasmids) do not carry AMR genes

R1-type plasmids (encoding Pfam01446) are often 2-3 kb in length and are typically comprised of a replication initiation protein and only two or three additional open reading frames encoding hypothetical proteins. R1-type plasmids constitute a small fraction of the plasmid dataset (n=16 plasmids, 13 isolates) and none of these carried AMR genes, suggesting that these plasmids are not yet involved in the acquisition and spread of AMR. Strains belonging to global clone 2 (GC2; largely represented by ST2) constitute over 90% of all *A. baumannii* genomes in GenBank, but R1 plasmids appear to be mainly associated with strains belonging to global clone 1 (GC1; largely represented by ST1, n=9/13 isolates) with only n=1 GC2 strain found with R1 plasmids (FIG 2).

**FIG 2.**
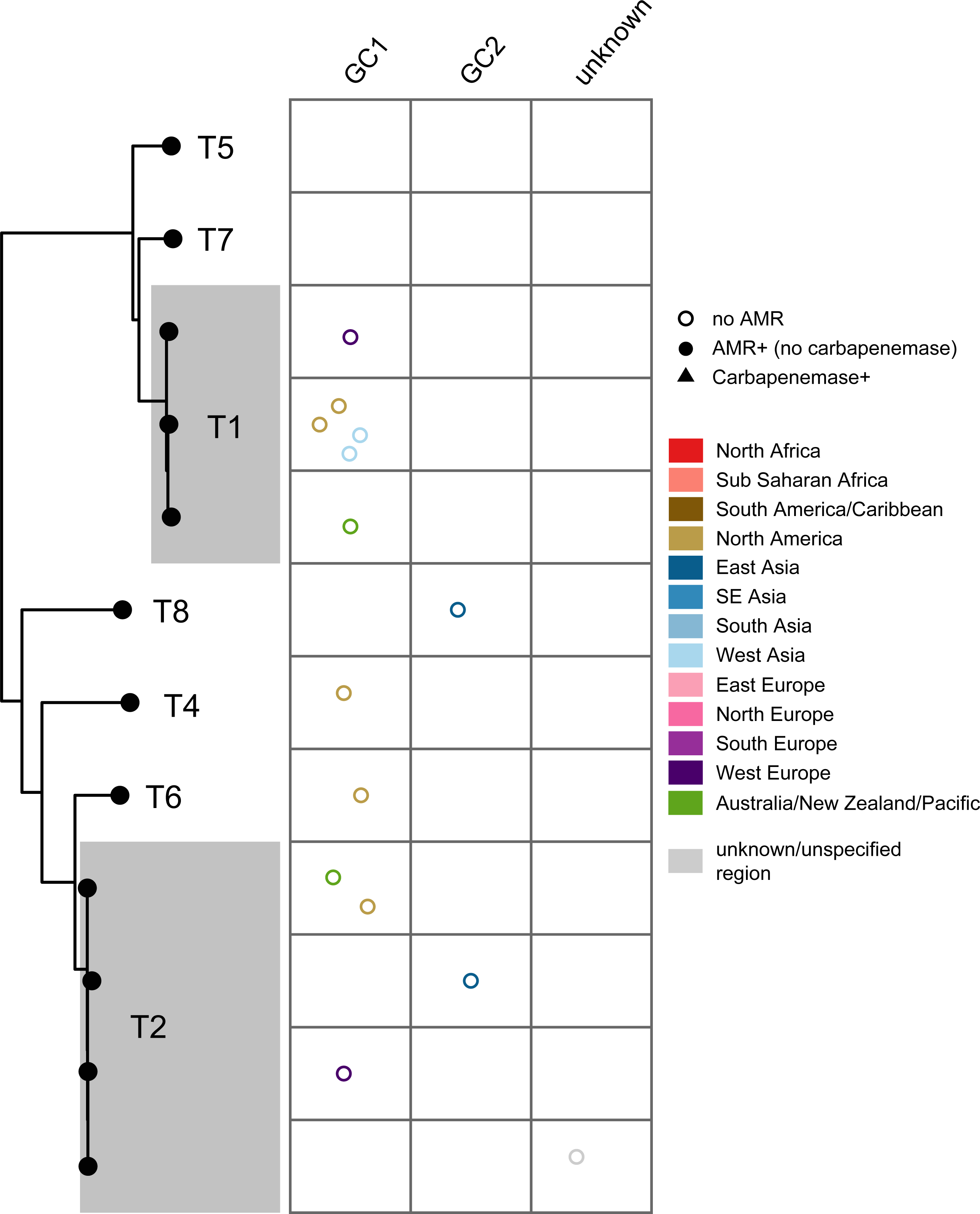
Phylogenetic relationship and distribution of antimicrobial resistance determinants in R1-type plasmids across major globally distributed sequence types and global clones. Plots show the plasmids linked with a particular plasmid type, with each data point corresponding to a unique plasmid, grouped by chromosomal sequence type/clone, and coloured by geographical region as shown in the Figure key. Empty circles (marked no AMR) indicate the absence of antimicrobial resistance gene.

### Rep_3 (R3) plasmids disseminate important AMR genes

R3 plasmids encoding the Rep_3-type plasmid replication proteins (Pfam01051) represent, by far, the most diverse *A. baumannii* plasmid group. This is due to several reasons, including the Rep/*rep* sequence divergence combined with their floating genetic structure arising from the presence of p*dif* modules (examples in FIG 3). Over half of the plasmids were typed as R3 (n=479/814 plasmids; 59%), and these were detected in at least 345 unique isolates (note, n=2 R3 type plasmids were unassigned to an isolate). Variants of R3-T1, T2 and T3 constitute the most abundant types and were collectively detected in n=224 plasmids. R3-type plasmids appear to be geographically dispersed, but some types appear to be limited to distinct regions (FIG 4). For example, some variants of R3-T1 plasmids that carry a carbapenemase appear to be limited to North America (e.g. pAB120 carrying *bla*_OXA-72_ from the US; GenBank accession number CP031446.1) or Europe (e.g. p1ABST78 from Italy; GenBank accession number AEOZ01000236.1 (FIG 4, Table S1 and Table S2), and may represent local plasmid circulation or expansions.

**FIG 3.**
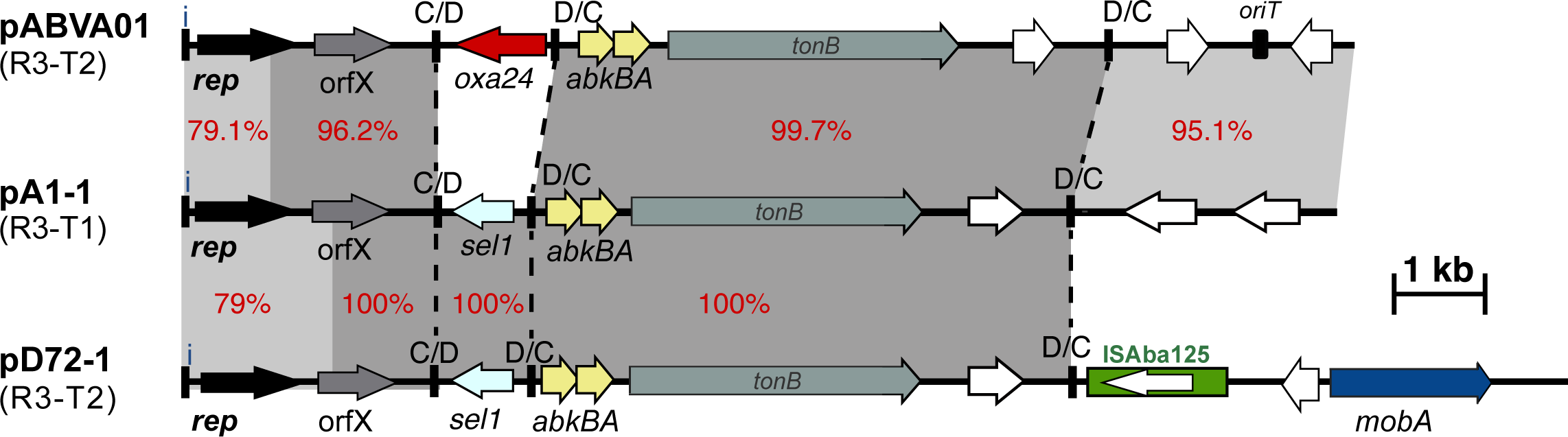
Schematic comparison of Rep_3 family (R3-type; Pfam01051) plasmid structures. Horizontal arrows show the length and orientation of genes with *rep* genes coloured black, resistance genes red, toxin/anti-toxins yellow and mobilisation genes blue. Green boxes indicate insertion sequences with their transposase shown inside the box. Small thick vertical bar marked with “i” indicate iterons. Dotted lines draw the show the boundaries of p*dif* modules. Other vertical bards marked with “C/D or D/C” indicate the location of p*dif* sites. Regions with significant DNA identities are shown using shades of grey with % identities also shown using red numbers.

**FIG 4.**
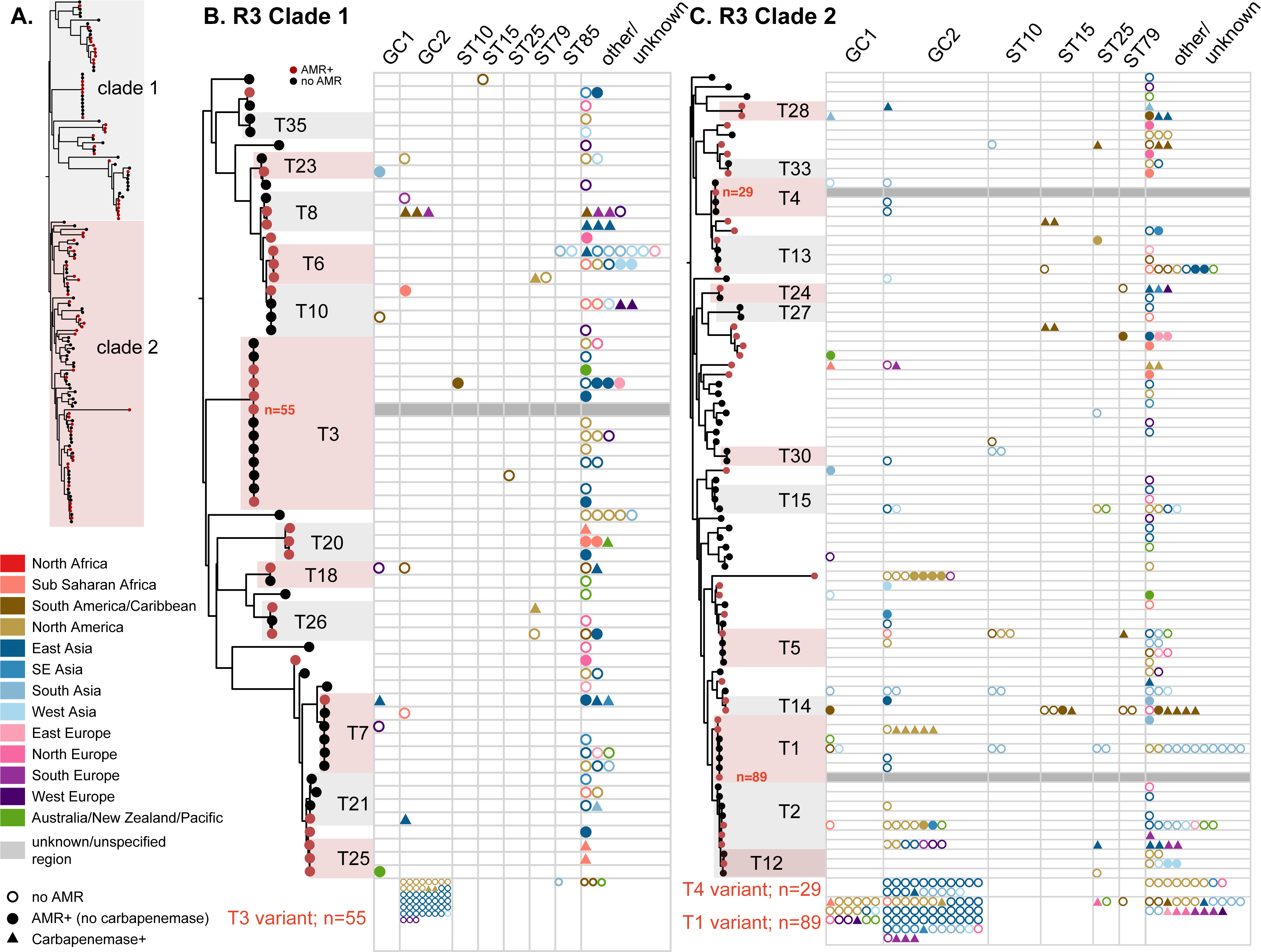
Phylogenetic relationship and distribution of antimicrobial resistance determinants in R3-type plasmids across major globally distributed sequence types and global clones. The overall phylogenetic tree is depicted in panel A), and clades 1 and 2 shown in greater resolution in panels B) and C), respectively. Nodes that are coloured red correspond to plasmid types where the presence of antimicrobial resistance genes is detected in at least one plasmid. Plots in panels B and C show the number of plasmids linked with a particular plasmid type; each data point corresponds to a unique plasmid, grouped by chromosomal sequence type/clone, and is coloured by geographical region as shown in the Figure key. Empty circles indicate plasmids with no AMR, triangles indicate plasmids with carbapenemases and filled circles represent plasmids with AMR (no carbapenemase). Data for three variants corresponding to R3-T3, R3-T4 and R3-T1 types are separately shown at due to spacing.

The updated R3 phylogeny generated from n=145 R3 *rep* nucleotide variants (n=78 distinct rep types) revealed two deep-branching clades containing n=61 and 84 *rep* types, with plasmids carrying AMR genes (including carbapenemases) dispersed across both clades. Approximately half of the R3 types were associated with plasmids without AMR (n=36/78 R3 types), while n=18 were linked only to plasmids that carry AMR, and n=24 (including the majority of the top 15 most common R3 types; see FIG 4) were associated with both AMR^+^ and AMR^-^ plasmids.

A quarter of R3-type plasmids (n=122/479; n=42 distinct R3 types collectively) were associated with AMR, and n=73 of these plasmids (from at least n=72 isolates) carried carbapenemases. Various AMR genes were detected on R3 plasmids; however, the *bla*_OXA-58_, *bla*_OXA-72,_ *bla*_OXA-24_ carbapenem resistance, *tet39* tetracycline resistance, *sul2* sulfonamide resistance and *msr-mph*(E) macrolide resistance genes were amongst the most abundant AMR genes found (Table S2). R3-type plasmids with AMR genes are carried by all major global clones including ST10, ST15, ST25, ST79 and ST85 strains, which collectively accounted for n=26/345 isolates carrying n=46 R3-type plasmids, but were predominantly detected in members of GC2 and GC1, accounting for n=150 and 26 isolates with R3-type plasmids, respectively (n=209/479 R3-type plasmids; see FIG 4 and Table 3).

Colistin is a last resort within our arsenal of antibiotics that largely remained effective against multi-drug resistant (MDR) *A. baumannii* ^18^. Here, the *mcr* colistin resistance gene was present in only n=5 isolates, all of which were linked to R3-type plasmids. These included 4 plasmids with the *mcr-4.3* gene carried by strains recovered in clinical samples (two strains isolated in each of China and the Czech Republic) on an R3-T22 plasmid (Table S2). The remaining plasmid, p8E072658, was recently described in an environmental isolate from recycled fibre pulp in a paper mill in Finland ^14^. This plasmid carries the novel *mcr-4.7* colistin resistance gene in a Tn*3*-family transposon and encodes two novel R3-type Reps (R3-T73 and R3-T74). Acquisition of the *mcr* plasmids is clinically significant given the importance of colistin in treatment of MDR *A. baumannii*, and while it currently appears to be uncommon, future monitoring of R3 plasmids with *mcr* may be warranted.

### The RP-type plasmids (encoding RepPriCT_1) disseminate aminoglycoside and carbapenem resistance genes

RP-type plasmids encoding RepPriCT_1 (Pfam03090) Rep have been reported in various *A. baumannii* strains and contribute to the emergence of MDR strains ^12,19–24^. Approximately one-fifth of the dataset (n=158 plasmids; 19.4%) were identified as RP-type. A significant portion of RP-type plasmids (n=64/158; 40.5%) carry at least one AMR gene highlighting the importance of this plasmid group in the acquisition and spread of AMR determinants. The majority of RP-type plasmids were typed as RP-T1 followed by RP-T2, accounting for n=130 and 22 plasmids (82.3% and 13.9%) respectively. The genetic structure of the predominant RP-T1 plasmid variant (pACICU2; GenBank accession number CP031382.1), is illustrated in Figure S1.

In fact, RP-T1 and RP-T2 plasmids accounted for all AMR^+^ RP-type plasmids. All n=49 RP-T1 AMR^+^ plasmids carried either *bla*_OXA-23_ (carbapenemase; n=31 RP-T1 plasmids) and/or *aphA6* (amikacin resistance; n=30 RP-T1); n=12 RP-T1 plasmids carried both. Other AMR genes detected in the RP-T1 plasmids included *sul1, dfrA7, aacA4, bla*_GES-11_*, strAB, aadA2, cmlA1, aadB,* and the *bla*_OXA-58_ carbapenemase (FIG 5 and Table S2). Moreover, it appears that variants of RP-T1 plasmids have similarly been acquired by all major globally distributed clones including members of GC1 and GC2, ST10, ST15, ST25, ST79 and ST622, recovered across all continents (FIG 5).

**FIG 5.**
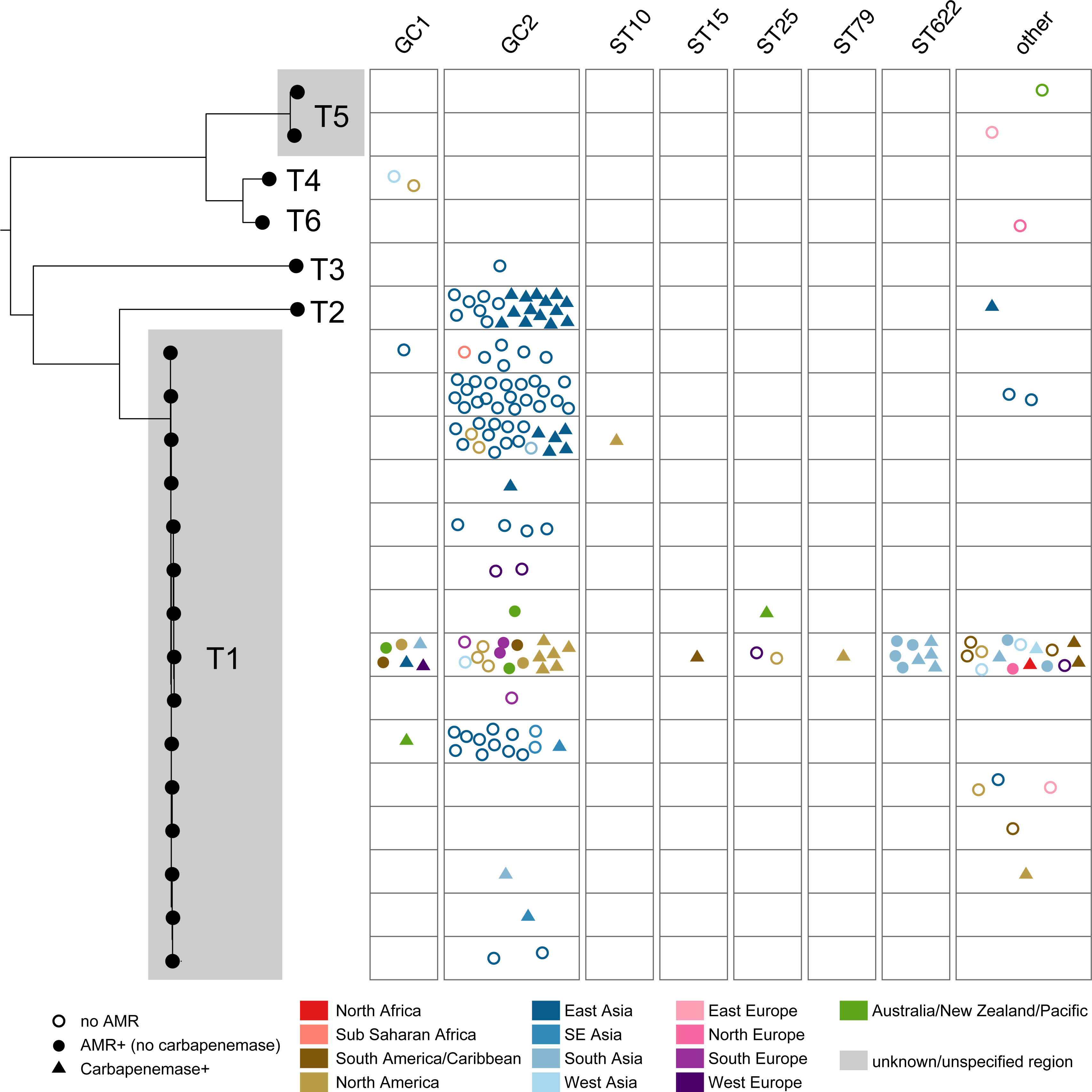
Phylogenetic relationship and distribution of antimicrobial resistance determinants in RP-type plasmids across major globally distributed sequence types and global clones. Plots show the number of plasmids linked with a particular plasmid type; each data point corresponds to a unique plasmid, grouped by chromosomal sequence type/clone, and is coloured by geographical region as shown in the Figure key. Empty circles indicate plasmids with no AMR, triangles indicate plasmids with carbapenemases and filled circles represent plasmids with AMR (no carbapenemase).

In contrast to the global distribution of RP-T1 plasmids, all n=22 RP-T2 plasmids were sequenced from isolates collected in East Asia (predominantly China, except for one plasmid with no AMR genes sequenced from an isolate in South Korea). Notably, n=15/22 plasmids contained a *bla*_OXA-23_ copy suggesting that RP-T2 plasmids with *bla*_OXA-23_ are circulating in China and have not yet been detected elsewhere. Plasmids corresponding to the remaining RP-types were generally small plasmids ranging in size from 4.5kb to 6.8kb (except for RP-T3; 52.5 kb) and carry no AMR genes. Interestingly, phylogenetic analysis of RP-type *rep* sequences (RepPriCT_1 family) revealed a clear separation of the smaller plasmids that lack AMR genes (RP-T4, RP-T5 and RP-T6) from the larger RP-T1, T2 and T3 plasmids (Figure X), suggesting distinct evolutionary trajectories that have likely influenced the accumulation of additional genes including those conferring AMR.

### Distribution of AMR genes in plasmids with no identifiable replication gene

We previously reported that a fraction of *A. baumannii* plasmids do not encode an identifiable replication initiation gene (i.e. 22.9%; n=142/621 plasmids) ^13^. Such plasmids might therefore use an alternative mechanism that does not involve a Rep to initiate replication or encode a novel Rep that is yet to be discovered. Here, n=161/814 plasmids did not encode an identifiable replication initiation gene. This *rep*-less group constitutes a set of highly diverse plasmids ranging in size from 4 kb to over 200 kb. Almost a third of these (n=52; 32.3%) appear to carry no AMR genes and range in size from 2.4 – 145.7 kb (Table 4). These plasmids are not discussed further as they lack AMR genes. The remaining n=109 plasmids (length range 3.8 kb to >200 kb) carry at least one AMR gene and constitute various plasmid variants. Some variants are associated with the carriage of clinically significant AMR genes, and include those related to pRAY*, large MPF_F_ conjugative plasmids such as pA297-3, and pNDM-BK01 (n=28, 31 and 8, respectively; accounting for 41.6% *rep*-less plasmids). These plasmids are further discussed below.

**Table 4.**
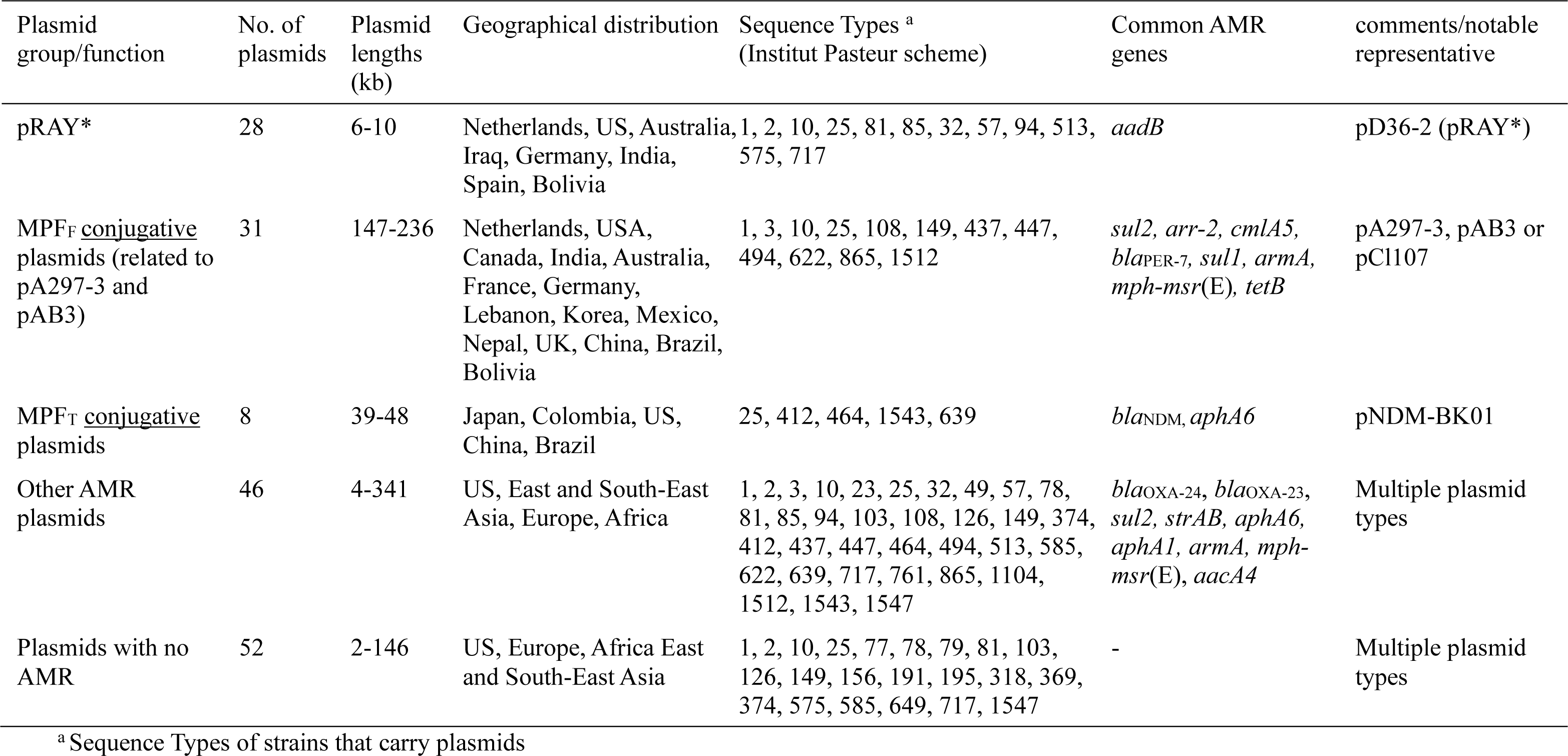
Summary of *rep*-less plasmids.

#### i) pRAY* – an important small plasmid spreading resistance to aminoglycosides

It has been shown that the small plasmid pRAY* and its variants play a role in the spread of the *aadB* gene conferring resistance to tobramycin, gentamicin and kanamycin, which are considered clinically significant antimicrobials ^25^. Although some variants did not carry an AMR gene (an example shown in FIG 6), we observed n=28 plasmids that were either identical or closely related to pRAY*, and most (n=25/28) carried *aadB.* These plasmids were found in strains assigned to at least 14 STs, including ST1, ST81, ST2, ST25 and ST85 (Table S3). The strains were also geographically diverse, indicating global dissemination of pRAY* plasmids. Moreover, all strains with pRAY* were recovered in clinical samples suggesting aminoglycoside selective pressures may play a role in driving their stable maintenance within clinical settings (Table S3).

**FIG 6.**
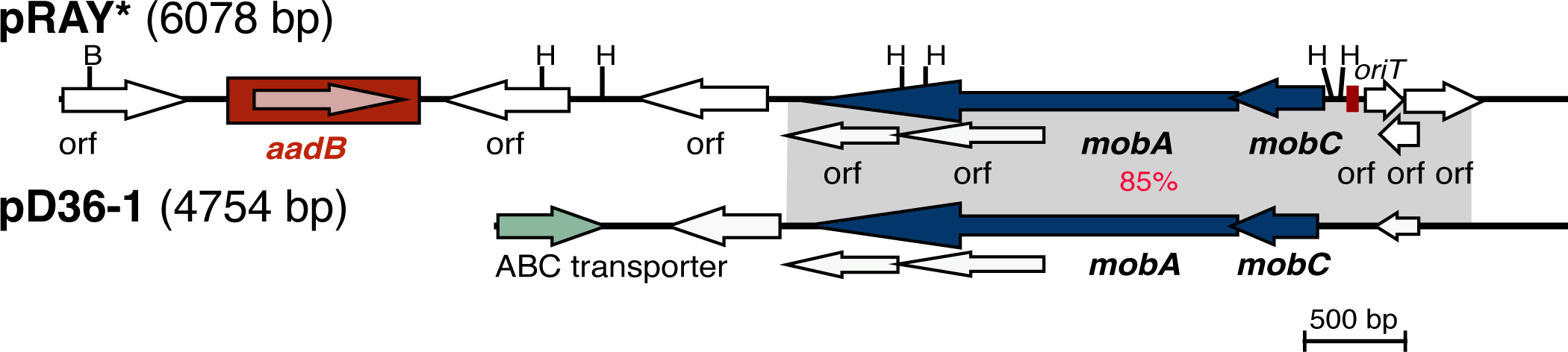
Linearised map of pD36-1 (cryptic) compared with pRAY* (pD36-2). Central horizontal lines indicate the plasmid backbones. Arrows represent the extent and orientation of genes, and the gene cassette is boxed. The grey shadings indicate regions with significant identity with the % identities indicated in red. Scale bar is shown. Drawn to scale from GenBank accession numbers CP012954 (pRAY*), and CP012953 (pD36-1).

#### ii) Spread of diverse AMR genes by conjugative plasmids encoding the MPF_F_ transfer system

This group constitutes a diverse set of n=31 large plasmids (146.7 – 236.2 kb in size) known to lack an identifiable *rep* gene. These plasmids were detected in at least 11 distinct STs with the highest count corresponding to ST622 (n=10) followed by ST25 (n=7) and were also present in members of the major global clones (e.g. ST1, ST10, ST25; Table S4). The 200.6 kb plasmid, pA297-3 (Table S4) is considered the representative as it has the most common backbone type and was one of the earliest described and shown to be conjugative ^11^. It carries *sul2* and *strAB,* conferring resistance to sulphonamide and streptomycin respectively. Most plasmids in this group (except p40288 and pR32_1; Table S4) carry a copy of *sul2.* Most members also carry *strAB* (n=28/31), *msr-mph*(E) (n=21), *bla*_PER-7_ (n=18), and *armA* (n=20) conferring resistance to streptomycin, macrolides, extended-spectrum β-lactamases (ESBLs), and aminoglycosides, respectively (Table S4). The latter, *armA,* encodes the 16S rRNA methylation protein that confers resistance to all aminoglycosides ^26^. Two plasmids, pPM193665_1 and pPM194122_1 (GenBank accession numbers CP050416 and CP050426, respectively) from strains recovered in India, also contain the *bla*_NDM_ metallo-ß-lactam carbapenem resistance gene.

#### iii) Conjugative plasmids encoding MPF_T_ transfer system

Though not very common, *bla*_NDM_ has now been reported in *A. baumannii* in several countries ^3,27–30^. In this study, we found n=8 plasmids with no identifiable *rep* gene that encode the MPF_T_ type conjugative transfer system ^31^ and carried the *bla*_NDM_ metallo-beta-lactam carbapenem resistance gene (Table 6). All these plasmids were found to be related to pNDM-BJ01 (GenBank accession number JQ001791.1), which was first reported in *Acinetobacter lowffii* and shown to be conjugative at a high frequency ^32^. These plasmids were carried by strains recovered in clinical, environmental (wastewater) and animal samples in different countries including China, Japan, US, Colombia, and Brazil showing their wide geographical distribution. They were found in various sequence types, of which only one (p1AR_0088; GenBank accession number CP027532.1; Table S2 and Table S5) was in a ST25 strain, which is an important globally distributed ST ^3,15,33–35^. Given the potential for accelerated resistance dissemination of resistance to a last-line antimicrobial and hence heightened therapeutic challenges, targeted surveillance of MPF_T_ type plasmids with *bla*_NDM_ may be warranted.

### Diverse plasmid types facilitating the spread of carbapenem resistance genes

Carbapenemases stand out as important AMR determinants as carbapenems are one of the last resort lines of defence in antimicrobial treatment ^3^. Here, we showed that various R3, RP and *rep*-less plasmids were associated with the spread of carbapenem resistance genes including *bla*o_XA-23_, *bla*_OXA-24_, *bla*_OXA-58_, and *bla*_NDM_. Carbapenemases were observed in 150 plasmids (18.4%), of which *bla*_OXA_-type genes were the most common carbapenemase type followed by *bla*_NDM_ (n=132, 11 and 7 plasmids with *bla*_OXA_ only, *bla*_NDM_ only and *bla*_OXA_ plus *bla*_NDM_ respectively). We detected twelve allelic variants of *bla*_OXA_-type genes; *bla*_OXA-23_ was the most prevalent (n=54), followed by *bla*_OXA-58_ (n=33, of which n=6 also carried *bla*_NDM-1_), *bla*_OXA-72_ (n=27), and *bla*_OXA-24_ (n=11). The prevalence of some of these alleles appear to be associated with distinct plasmid types. For example, n=46/54 *bla*_OXA-23_ plasmids were typed as RP-T1 or RP-T2, while R3-type plasmids appear to play a key role in the dissemination of *bla*_OXA-58_ (i.e. n=28/33 plasmids with *bla*_OXA-58_), *bla*_OXA-72_ (n=21/27), and *bla*_OXA-24_ (n=10/11). Notably, except for *bla*_OXA-23_, which was abundant in ST2 isolates (n=33/54 *bla*_OXA-23_ plasmids), there was no clear correlation of carbapenemase-carrying plasmids with particular sequence types (ST), indicating widespread distribution of plasmids between various *A. baumannii* clones.

The *bla*_NDM_ carbapenem resistance gene is clinically significant in all Gram-negative bacteria, especially Enterobacterales as its rapid spread among different bacterial species worldwide has become a serious threat to public health ^36^. A single allelic variant, *bla*_NDM-1_, was observed in this dataset and detected in n=13 plasmids including n=8 pNDM-BJ01-type variants, n=2 R3-type n=2 pA297-3-type, and a novel plasmid pCCBH31258 (GenBank accession number CP101888). The presence of *bla*_NDM_ on conjugative plasmids in *A. baumannii* is significant as it highlights the potential for the rapid transmission of this important carbapenemase via horizontal gene transfer.

### Opportunities and limitations

Advances in whole genome sequencing technologies combined with the rapid accumulation of genome data in publicly available databases such as GenBank has provided a valuable opportunity to gain genomic insights towards the circulation of AMR genes in critical pathogens such as *A. baumannii* and, more importantly, MGEs that disseminate AMR. However, this unique opportunity is associated with important caveats given that publicly available genome sequences are largely geographically skewed for several reasons, including the lack of technology, financial support and expertise in developing countries. Currently, the bulk of genome sequence data in GenBank has been sequenced from isolates collected in the US, China and Australia; these countries accounted for approx. ∼50% of the dataset in this study. The geographical skew of genome sequence data makes it difficult to gain comprehensive insights into population structure, MGEs and AMR genes circulating in other parts of the world e.g. Africa and the Middle East. Moreover, genomes of environmental *A. baumannii* strains are also very limited (FIG 1), making it difficult to understand the distribution of AMR genes and MGEs including plasmids in the environment. Consequently, the scarcity of environmental genome data also limits the study of how plasmids, particularly those with AMR, circulate between the strains from clinical and environmental origins.

## CONCLUSIONS

Traditionally, *A. baumannii* has been characterized as an organism that primarily acquires AMR genes through large chromosomal islands. However, this definition is changing as more plasmids that carry important AMR genes are being characterised. This study also highlights the pivotal role of various plasmid types, particularly certain families in the dissemination of clinically important AMR genes within this pathogen. We showed that many RP-T1, R3-types (e.g. RP-T1 and R3-T2) and rep-less pNDM plasmids can spread various carbapenem resistance genes. These are of particular concern as AMR genes conferring resistance to carbapenems are often considered as the last line of defence in treatment. Furthermore, these plasmids typically carry additional AMR genes conferring resistance to multiple antimicrobials, which further compounds treatment management and the threat posed by *A. baumannii*.

Although the plasmid repertoire of *A. baumannii* exhibits remarkable diversity, this investigation highlights the profound significance of specific plasmid families in harboring and disseminating AMR genes. The findings from this study provide new insights into which plasmid types are over-represented among those that disseminate AMR and may be flagged as targets for focused AMR surveillance. Finally, this study showed that, in the ongoing battle against antibiotic resistance in *A. baumannii,* its plasmids play a significant role in exacerbating the crisis. Their ability to transfer AMR genes across different sequence types, coupled with the bacterium’s adaptability, poses a formidable challenge to healthcare systems worldwide.

## MATERIALS AND METHODS

### Plasmid sequence data

A local database of complete *A. baumannii* plasmids that were publicly available as of mid-August 2022 was generated. Our local database included plasmid sequences of i) 355 complete genomes out of 450 complete *A. baumannii* genomes (i.e. n=95 entries with no plasmids) sourced from GenBank (https://www.ncbi.nlm.nih.gov/genome/browse/#!/prokaryotes/403/); labelled as ‘Complete genome project’ as data source in Table S1 and Table S2) and ii) an additional 92 genomes/unique strains (released between February 2021 and mid-August 2022) captured in RefSeq (https://www.ncbi.nlm.nih.gov/refseq/). The latter included n=29 genomes were sourced from Whole Genome Shotgun projects (labelled as ‘WGS’ in Table SX) and n=63 unique strains that were not linked to a genome project (i.e. direct plasmid submission to GenBank; labelled as ‘GenBank non-redundant db’ in Table S2). This resulted in the curation of our final dataset consisted of 814 non-redundant plasmid entries corresponding to at least 440 unique isolates (n=355 isolates from Complete genome projects, n=63 from GenBank non-redundant database, and n=29 from WGS). Of the 814 plasmid entries, n=621 were those we previously used to develop the *Acinetobacter* Plasmid Typing scheme ^13^. All supporting data and protocols have been provided within the article or through supplementary data files. The online version of this article has four supplementary tables and three supplementary figures.

### Bioinformatics and sequence analysis

The chromosomal sequences associated each plasmid were found by exporting the BioSample accession numbers using the RefSeq https://www.ncbi.nlm.nih.gov/refseq/ followed by the curation of a list of chromosomal GenBank accession numbers and downloading the sequence data through Entrez Programming Utilities (E-utilities; https://www.ncbi.nlm.nih.gov/books/NBK25501/). Chromosomal sequences were used to determine the Multi-locus Sequence Types (MLSTs) using the *mlst* v.2.0 software (https://github.com/tseemann/mlst). Standalone BLAST (https://ftp.ncbi.nlm.nih.gov/blast/executables/LATEST/) was used for plasmid sequence comparisons within the *rep*-less plasmid group and assign ‘related known plasmid’ variants as labelled in Table S2. The SnapGene® (V.6.0.5) software was used to examine the structure of individual plasmids. The plasmids were screen for AMR genes using Abricate v1.0.1 (available at https://github.com/tseemann/abricate) using the ResFinder v.2.1 database (available under https://cge.cbs.dtu.dk/services/ResFinder/). Data visualisation was performed using the ggplot2 package (https://ggplot2.tidyverse.org/) in R (v1.1.456) and Adobe Illustrator (V23.0.3).

### Clustering and phylogenetic analysis of the *rep/*Rep sequence data

*rep/*Rep sequences were extracted from the novel plasmid entries using the SRST2 software ^37^ followed by manual curation. Clusters comprising *rep* sequences at >95% nucleotide identity were derived using CD-HIT Suite (https://github.com/weizhongli/cdhit) ^38^, as previously described ^13^. The *rep* nucleotide sequences were separately aligned for each of the Rep families using MUSCLE v3.8.31 ^39^. Phylogenies were generated using the aligned *rep* sequences as input into RAxML v8.2.9 run five times with the generalised time-reversible (GTR) model and a Gamma distribution. The final trees with the highest likelihoods were selected, visualised in FigTree v1.4.4 (http://tree.bio.ed.ac.uk/), and annotated with the plotTree code (https://github.com/katholt/plotTree) in R v1.1.456.

## ACKNOWLEDGEMENTS

M.M.C.L. and M.H., conceptualisation, formal analysis, methodology. M.H. writing – original draft. M.M.C.L. and M.H. writing – review & editing. MH, funding.

M.M.C.L. is supported by an Australian National Health and Medical Research Council Investigator Grant (APP2009163). M.H. is supported by an Australian Research Council DECRA fellowship (DE200100111).

The authors declare no Conflict of Interest.

## FIGURE LEGENDS

**Supplementary FIG 1** Genetic structure of pACICU2 representing plasmids that encode RP-T1 replication initiation protein. Arrows indicate the extent and orientation of genes and open reading frames with red showing resistance genes, black *rep,* and flax transfer genes. Boxes coloured green indicate ISAba125 copies.

**Supplementary FIG 2** Circular map of pA297-3 drawn to scale from GenBank accession number KU744946. Arrows represent the orientation and extent of genes and open reading frames. Open reading frames with no predicted function are white and antimicrobial resistance genes are coloured red. Insertion sequences (IS) are shown with filled boxes coloured different shades of green. Arrows coloured flax represent the *tra* genes, which are involved in plasmid transfer. Gray arrows represent genes/orfs involved in DNA metabolism.

## SUPPLEMENTARY MATERIAL

**Table S1:** Available at https://doi.org/10.6084/m9.figshare.24076776

**Table S2:** Available at https://doi.org/10.6084/m9.figshare.24076779

**TABLE S3.** Properties of strains carrying pRAY* and its variants.

| Plasmid name | Strain name | Length (bp) | ST | Year | Isolation source | Country | Accession number |
| --- | --- | --- | --- | --- | --- | --- | --- |
| pA297-1 | A297 (RUH875) | 6078 | 1 | 1984 | nr | Netherlands | KU869529 |
| pD36-2 | D36 | 6078 | 81 | 2008 | wound | Australia | CP012954 |
| pMRSN3527-6 | MRSN 3527 | 6068 | 81 | 2011 | wound | USA | CM003318 |
| pRAY*-v1 | C2 | 6078 | 2 | 2007 | nr | Australia | JF343536 |
| p3ZQ2 | ZQ2 | 6078 | 2 | 2016 | sputum | Iraq | CM009648 |
| pABLAC2 | LAC-4 | 6076 | 10 | 1997 | HO <sup>a</sup> | USA | CP007714 |
| pD46-1 | D46 | 6078 | 25 | 2010 | UTI <sup>b</sup> | Australia | CP048132 |
| pNaval18-6.1 | Naval-18 | 6078 | 25 | 2006 | nr | USA | AFDA02 <sup>c</sup> |
| pR32_3 | Nord4-2 | 11378 | 25 | 2018 | nr | Germany | CP091597 |
| p6ACN21 | ACN21 | 9909 | 85 | 2018 | blood | India | CP038647 |
| p4ACN21 | ACN21 | 7396 | 85 | 2018 | blood | India | CP038649 |
| p3ACN21 | ACN21 | 6944 | 85 | 2018 | blood | India | CP038650 |
| p2ACN21 | ACN21 | 5844 | 85 | 2018 | blood | India | CP038651 |
| p1ACN21 | ACN21 | 5734 | 85 | 2018 | blood | India | CP038652 |
| pAbBAS-1.2 | AbBAS-1 | 6078 | 85 | 2019 | clinical | Spain | CP065394 |
| pMRSN4106-6 | MRSN 4106 | 6078 | 94 | 2011 | wound | USA | CM003315 |
| pMRSN3942-6 | MRSN 3942 | 6078 | 94 | 2011 | wound | USA | CM003319 |
| pMRSN3405-6 | MRSN 3405 | 6078 | 94 | 2011 | wound | USA | CM003320 |
| pJ9-1 | J9 | 6078 | 49 | 1999 | clinical | Australia | CP041588 |
| p1AR_0070 | AR_0070 | 6078 | 32 | nr | clinical | USA | CP027181 |
| p1AR_0052 | AR_0052 | 6078 | 32 | nr | clinical | USA | CP027187 |
| p2ZQ10 | ZQ10 | 6078 | 575 | 2016 | CSF <sup>d</sup> | Iraq | CM009031 |
| p3ZQ9 | ZQ9 | 6133 | 575 | 2016 | blood | Iraq | CM009085 |
| p1ZQ3 | ZQ3 | 6078 | 717 | 2016 | blood | Iraq | CM009028 |
| p2ZQ8 | ZQ8 | 6078 | 513 | 2016 | blood | Iraq | CM009034 |
| p1FDAARGOS_533 | FDAARGOS_533 | 6078 | 57 | 2016 | sputum | USA | CP033770 |
| pRAY*-v2 | E7 | 8433 | novel | 2008 | blood | Australia | JX076770 |
| pMC23.3 | MC23 | 6078 | novel | 2016 | urine | Bolivia | MK531539 |

<sup>a</sup> hospital outbreak – site not recorded.
<sup>b</sup> urinary tract infection (UTS)
<sup>c</sup> complete GenBank accession number AFDA02000006
<sup>d</sup> cerebrospinal fluid (CSF)

**TABLE S4.** Distribution of antimicrobial resistance genes in large conjugative plasmid related to pA297-3.

| Plasmid name | Length (bp) | ST | Year | Isolation source | Country | Antimicrobial resistance genes | Accession number |
| --- | --- | --- | --- | --- | --- | --- | --- |
| pA297-3 | 200633 | 1 | 1984 | UTI | Netherlands | *sul2, strAB* | KU744946 |
| pOIFC137-122 | 122461 | 3 | 2003 | nr<sup>a</sup> | USA | *strAB, sul2* | AFDK01000004 |
| pOIFC109-122 | 122469 | 3 | 2003 | nr | USA | *strAB, sul2* | ALAL01000013 |
| pAB3 | 148955 | 437 | 2014 | clinical | Canada | *sul2* | CP012005 |
| pAB04-1 | 169023 | 10 | 2012 | blood | Canada | *aph(3'')-Ib, strAB, sul2, arr-2, cmlA5, bla<sub>PER-7</sub>, sul1, armA, mph-msr(E), tetB* | CP012007 |
| pPM193665_1 | 150385 | 10 | 2019 | Pus | India | *mph-msr(E), armA, sul1, cmlA5, arr-2, sul2, strAB, ble<sub>MBL</sub>, bla<sub>NDM</sub>, tetB* | CP050416 |
| pPM194122_1 | 150385 | 10 | 2019 | BAL<sup>b</sup> | India | *mph-msr(E), armA, sul1, cmlA5, arr-2, sul2, strAB, ble<sub>MBL</sub>, bla<sub>NDM</sub>, tetB* | CP050426 |
| pOIFC143-128 | 127633 | 25 | 2003 | nr | USA | *strAB, sul2* | AFDL01000008 |
| pD4 | 132632 | 25 | 2006 | wound | Australia | *sul2, strAB* | CP048851 |
| pD46-4 | 208004 | 25 | 2010 | UTI | Australia | *tetB, sul2, mph-msr(E), strAB* | CP048135 |
| p40288 | 145711 | 25 | 2015 | UTI | France | - | CP077802 |
| pR32_1 | 117234 | 25 | 2018 | nr | Germany | - | CP091598 |
| pCL107 | 198716 | 25 | 2012 | UTI | Lebanon | *sul2, tetB, strAB, aacC2* | CP098522 |
| pNaval18-131 | 130660 | 25 | 2006 | nr | USA | *sul2, strAB* | AFDA02000009 |
| pHWBA8_1 | 195838 | 25 | 2013 | sputum | Korea | *armA, mph-msr(E), tetB, aac(6')-lan, aac(3)-lle, sul2, arr-2, cmlA5, sul1, bla<sub>PER-7</sub>* | CP020596 |
| pAba7804b | 170420 | 25 | 2006 | BAL | Mexico | *sul2, tetB, strAB* | CP022285 |
| p2AR_0088 | 146698 | 25 | nr | clinical | USA | *sul2, tetB, strAB, aac(3)-lle, aac(6')-lan* | CP027531 |
| p3P7774 | 202283 | 25 | 2018 | Pus | India | *tetB, mph-msr(E), armA, bla<sub>PER-7</sub>, sul1, cmlA5, arr-2, sul2, strAB, aac(6')-lan* | CP040260 |
| pVB82_1 | 215278 | 25 | 2019 | blood | India | *aac(6')-lan, bla<sub>OXA-23</sub>, strAB, sul2, tetB, mph-msr(E), armA, sul1, bla<sub>PER-7</sub>, cmlA5, arr-2* | CP050386 |
| p2VB16141 | 189343 | 622 | 2019 | blood | India | *strAB, sul2, arr-2, cmlA5, bla<sub>PER-7</sub>, sul1, armA, mph-msr(E)* | CP040051 |
| pIOMTU433 | 189354 | 622 | 2013 | clinical | Nepal | *mph-msr(E), armA, bla<sub>PER-7</sub>, sul1, cmlA5, arr-2, sul2, strAB* | AP014650 |
| p1KSK6 | 218105 | 622 | 2020 | respiratory | India | *mph-msr(E), armA, bla<sub>PER-7</sub>, sul1, cmlA1, arr-3, ant(3'')-la, sul2, strAB,* | CP072271 |
| p1KSK7 | 218105 | 622 | 2020 | respiratory | India | *mph-msr(E), armA, bla<sub>PER-7</sub>, sul1, cmlA1, arr-3, ant(3'')-la, sul2, strAB,* | CP072276 |
| p1KSK10 | 218105 | 622 | 2020 | respiratory | India | *mph-msr(E), armA, bla<sub>PER-7</sub>, sul1, cmlA1, arr-3, ant(3'')-la, sul2, strAB,* | CP072281 |
| p1KSK11 | 218105 | 622 | 2020 | respiratory | India | *mph-msr(E), armA, bla<sub>PER-7</sub>, sul1, cmlA1, arr-3, ant(3'')-la, sul2, strAB,* | CP072286 |
| p1KSK18 | 218105 | 622 | 2020 | respiratory | India | *mph-msr(E), armA, bla<sub>PER-7</sub>, sul1, cmlA1, arr-3, ant(3'')-la, sul2, strAB,* | CP072291 |
| p1KSK19 | 218105 | 622 | 2020 | respiratory | India | *mph-msr(E), armA, bla<sub>PER-7</sub>, sul1, cmlA1, arr-3, ant(3'')-la, sul2, strAB,* | CP072296 |
| p1KSK20 | 218105 | 622 | 2020 | respiratory | India | *mph-msr(E), armA, bla<sub>PER-7</sub>, sul1, cmlA1, arr-3, ant(3'')-la, sul2, strAB,* | CP072301 |
| p1KSK2 | 218105 | 622 | 2020 | respiratory | India | *mph-msr(E), armA, bla<sub>PER-7</sub>, sul1, cmlA1, arr-3, ant(3'')-la, sul2, strAB,* | CP072399 |
| pNCTC7364 | 148956 | 494 | 2014 | nr | UK | *sul2* | LT605060 |
| pB11911 | 216780 | 149 | 2014 | blood | India | *strAB, sul2, arr-2, cmlA5, bla<sub>PER-7</sub>, sul1, armA, mph-msr(E)* | CP021344 |
| p3VB35179 | 236166 | 1512 | 2018 | blood | India | *strAB, sul2, arr-2, cmlA5, bla<sub>PER-7</sub>, sul1, armA, mph-msr(E), tetB* | CP040054 |
| pA1429c | 205113 | 108 | 2010 | secretion | China | *bla<sub>TEM-1</sub>, aac(3)-lle, aac(6')-lan, strAB, tetB, sul2* | CP046899 |
| pPM194229_1 | 226394 | 447 | 2019 | BAL | India | *mph-msr(E), armA, sul1, bla<sub>PER-7</sub>, cmlA5, arr-2, strAB, sul2, tetB* | CP050433 |
| p1KSK1 | 218105 | 865 | 2020 | respiratory | India | *mph-msr(E), armA, sul1, bla<sub>PER-7</sub>, cmlA1, arr-3, ant(3'')-la, strAB,* | CP072123 |
| pAb45063_b | 183767 | nk | nr | nr | Brazil | *sul2, strAB* | MK323043 |
| pMC1.1 | 184770 | nk | 2015 | catheter | Bolivia | *aacC5, aac(6')-lan, strAB, tetB, sul2* | MK531536 |
| pMC75.1 | 150158 | nk | 2016 | ulcer | Bolivia | *strA, aph(6), sul2* | MK531540 |

<sup>a</sup> not recorded
<sup>a</sup> Broncho-alveolar lavage (BAL)

**TABLE S5.** Properties of plasmids encoding the MPF_T_ transfer system that carry *bla*_NDM_.

| Plasmid name | Length (bp) | Isolation source | Country | Year | ST | Antimicrobial resistance genes | Accession number |
| --- | --- | --- | --- | --- | --- | --- | --- |
| pOCU_Ac16a_2 | 41087 | tracheal aspirate | Japan | 2015 | 412 | *bla*<sub>NDM-1</sub>, *bleMBL*, *aphA6* | AP023079 |
| p6200-47.274kb | 47274 | bodily fluid | Colombia | 2012 | 464 | *bla*<sub>NDM-1</sub>, *bleMBL*, *aphA6* | CP010399 |
| pNDM-0285 | 39359 | wastewater | USA | 2016 | 1543 | *bla*<sub>NDM-1</sub>, *bleMBL*, *aphA6* | CP026127 |
| p1AR_0088 | 41087 | clinical | USA | <2013 | 25 | *bla*<sub>NDM-1</sub>, *bleMBL*, *aphA6* | CP027532 |
| pAbNDM-1 | 48368 | feces | China | <2013 | 639 | *bla*<sub>NDM-1</sub>, *bleMBL*, *aphA6* | JN377410 |
| pNDM-AB | 47098 | pig lung | China | <2013 | novel | *bla*<sub>NDM-1</sub>, *aphA6*, *mph-msr(E)* | KC503911 |
| Piec383 | 47283 | blood | Brazil | 2014 | novel | *bla*<sub>NDM-1</sub> | MK053932 |
| pAB17 | 41087 | - | Brazil | <2020 | novel | *bla*<sub>NDM-1</sub>, *aphA6* | MT002974 |

