## Supplementary figures and images for "Examining the role of *Acinetobacter baumannii* Plasmid Types in Disseminating Antimicrobial Resistance"

### FigS1-pAb-G7-2-Map_12042023.pdf

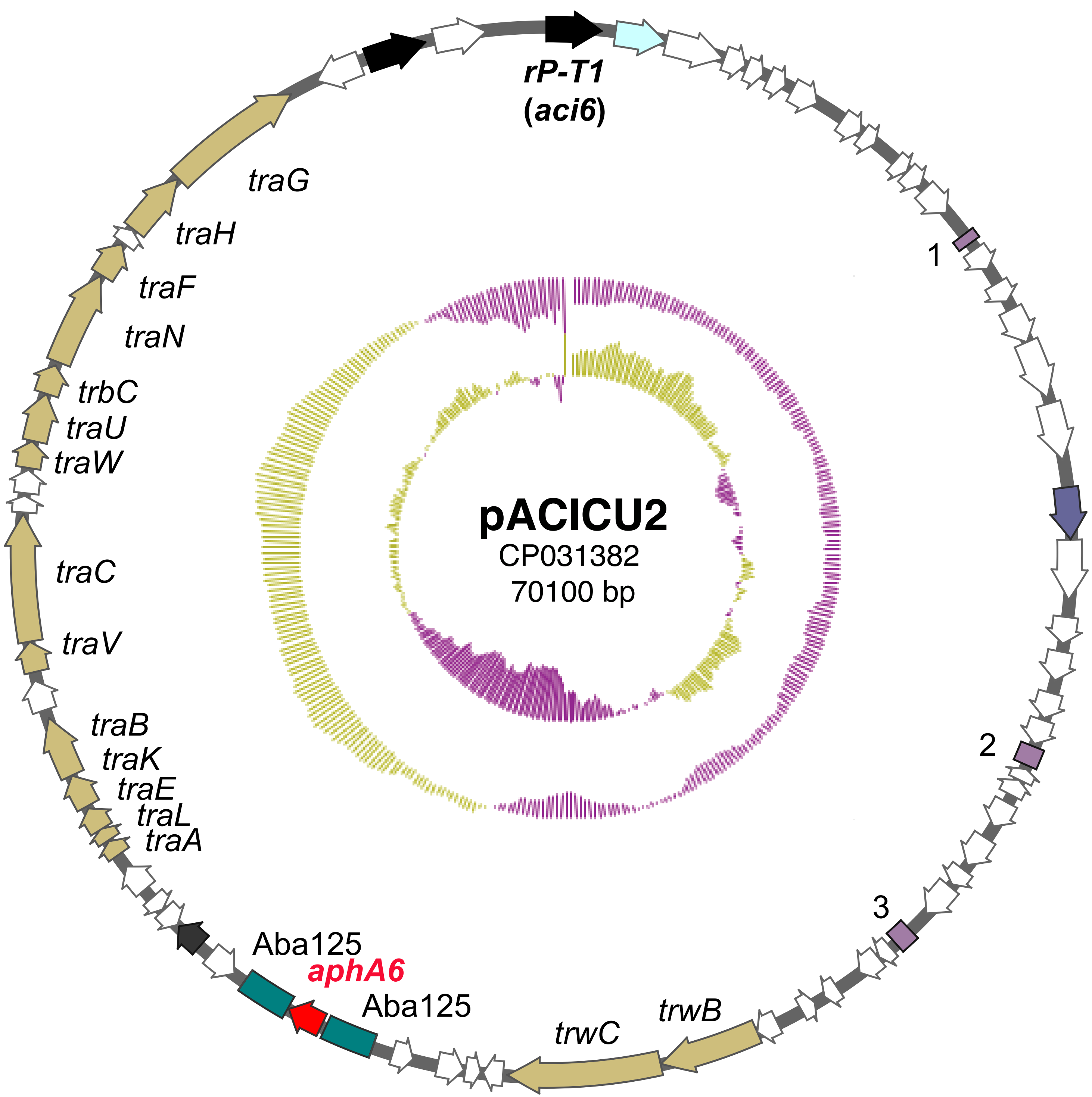

### FigS2_pA293-map.pdf

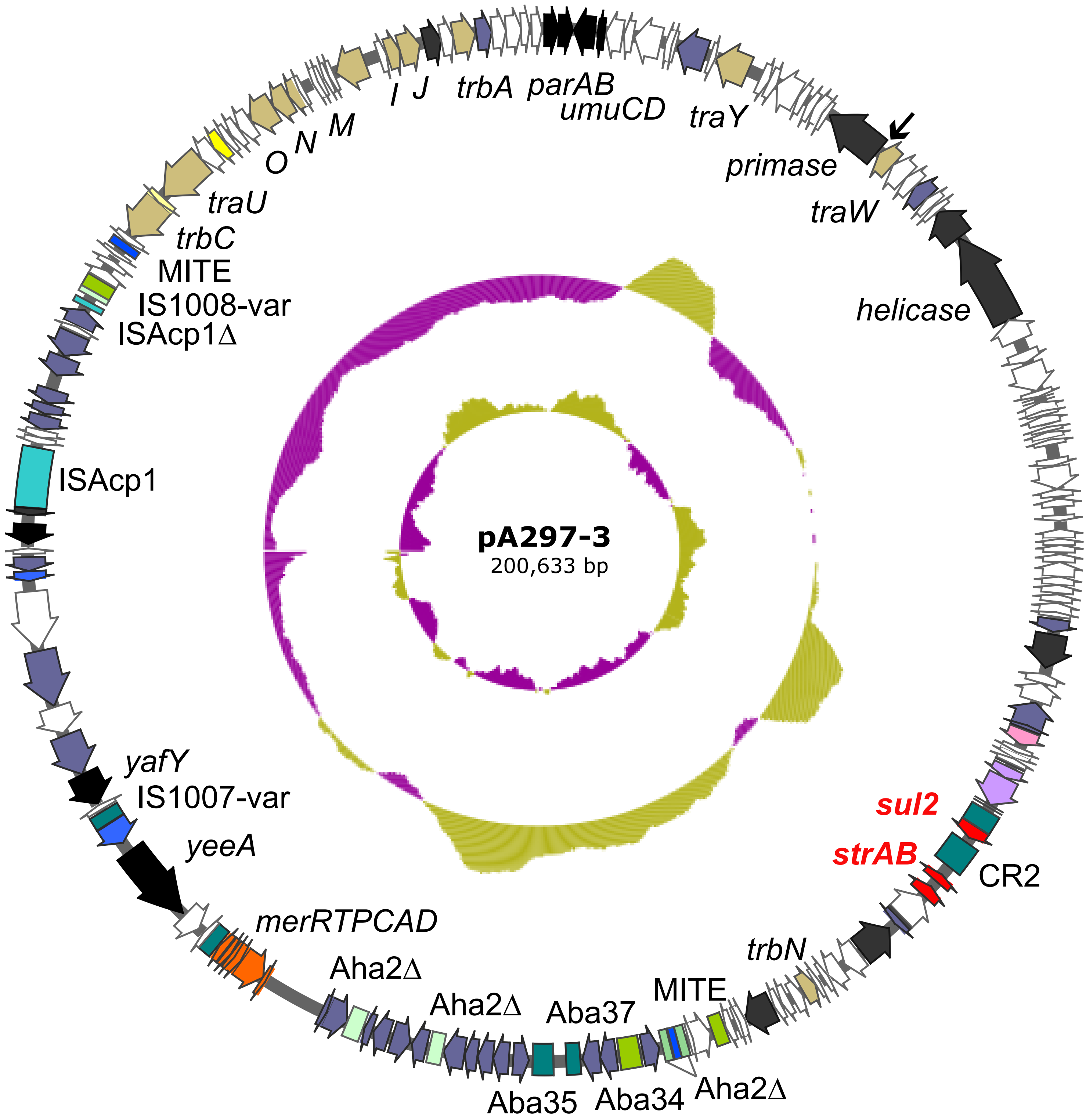
